# Murine gammaherpesvirus 68 ORF45 stimulates B2 retrotransposon and pre-tRNA activation in a manner dependent on mitogen-activated protein kinase (MAPK) signaling

**DOI:** 10.1101/2022.07.09.499448

**Authors:** Azra Lari, Britt A. Glaunsinger

## Abstract

RNA polymerase III (RNAPIII) transcribes a variety of noncoding RNAs, including transfer RNA (tRNA) and the B2 family of short interspersed nuclear elements (SINEs). B2 SINEs are noncoding retrotransposons that possess tRNA-like promoters and are normally silenced in healthy somatic tissue. Infection with the murine gammaherpesvirus MHV68 induces transcription of both SINEs and tRNAs, in part through the activity of the viral protein kinase encoded by ORF36. Here, we identify the conserved MHV68 tegument protein ORF45 as an additional activator of these RNAPIII loci. MHV68 ORF45 and ORF36 form a complex, resulting in an additive induction RNAPIII and increased ORF45 expression. ORF45-induced RNAPIII transcription is dependent on its activation of the extracellular signal-regulated kinase (ERK) mitogen-activated protein kinase (MAPK) signaling pathway, which in turn increases the abundance of the RNAPIII transcription factor Brf1. Other viral and non-viral activators of MAPK/ERK signaling also increase the levels of Brf1 protein, B2 SINE RNA and tRNA, suggesting that this is a common strategy to increase RNAPIII activity.

**Importance:** Gammaherpesviral infection alters the gene expression landscape of a host cell, including through induction of non-coding RNAs transcribed by RNA polymerase III (RNAPIII). Among these are a class of repetitive genes known as retrotransposons, which are normally silenced elements that can copy and spread throughout the genome, and transfer RNAs (tRNAs), which are fundamental components of protein translation machinery. How these loci are activated during infection is not well understood. Here, we identify ORF45 from the model murine gammaherpesvirus MHV68 as a novel activator of RNAPIII transcription. To do so, it engages the MAPK/ERK signaling pathway, which is a central regulator of cellular response to environmental stimuli. Activation of this pathway leads to upregulation of a key factor required for RNAPIII activity, Brf1. These findings expand our understanding of the regulation and dysregulation of RNAPIII transcription and highlight how viral co-option of key signaling pathways can impact host gene expression.

## Introduction

Mammalian genomes retain a remarkably high number of repetitive sequences derived from transposable elements. Retrotransposons are the largest group among these ancient invasive DNA elements and are defined by the ability to mobilize throughout the genome via RNA intermediates (1, 2). Approximately 10% of the mammalian genome consists of a class of retrotransposons known as short interspersed nuclear elements (SINEs) (1, 3, 4). SINEs are short (<500bp), non-autonomous, noncoding retrotransposons transcribed by RNA Polymerase III (RNAPIII). Both human and murine genomes harbor SINE families evolutionarily derived from endogenous RNAPIII transcripts. Alu elements, the predominant family of SINEs in the human genome, and B1 elements in the murine genome are both derived from the 7SL RNA, which is the RNA component of the signal recognition particle (5–7). The unrelated murine B2 SINE family is derived from transfer RNA (tRNA) (8). Despite their independent origins, human and murine SINEs are ubiquitously present across genomes and are often localized proximal to or within protein encoding genes (9–11). As such, SINE sequences are known to act as functional enhancers or mobile promoters for proximal genes and can impact the processing, localization and decay of mRNA transcripts when present as embedded elements (3, 12–27).

Transcription of SINEs is typically repressed in healthy somatic cells in part due to the maintenance of repressive tri-methylation of lysine 9 on histone H3 (H3K9) marks and CpG methylation of SINE DNA (28–32). However, in response to cellular stresses such as chemical treatment, heat stress, or viral infection, SINEs are de-repressed and robustly transcribed into SINE non-coding RNAs (ncRNAs) (33–39). Both human and murine SINEs are actively transcribed during infection with several DNA viruses including herpes simplex virus (HSV-1), adenovirus type 5, minute virus of mice (MVM), simian virus 40 (SV40), and murine gammaherpesvirus 68 (MHV68), which is related to the human gammaherpesviruses Kaposi’s sarcoma-associated herpesvirus (KSHV) and Epstein-Barr virus (EBV) (34, 37, 40–43).

Upregulation of RNAPIII transcripts during viral infection is not limited to SINEs. Many DNA viruses encode their own RNAPIII transcribed genes whose expression is stimulated by enhanced RNAPIII activity during infection (40, 43, 44). Infection with DNA viruses can also enhance transcription and abundance of host premature tRNAs (pre-tRNAs); recent genome wide studies have identified a global increase in pre-tRNA levels during MHV68 and HSV-1 infection (45, 46).

SINE ncRNAs induced during cellular stress have a variety of documented functions. For example, both Alu and B2 SINE ncRNAs interact directly with RNA Polymerase II (RNAPII) to repress gene expression from individual promoters (47–49). During MHV68 infection, cytoplasmic B2 SINE ncRNAs can activate NF-κB innate immune signaling pathways leading to increased viral replication (39). Constitutive expression of Alu ncRNAs has also been linked to pathogenic outcomes such as age-related macular degeneration (AMD) through activation of the inflammasome, and increased epithelial-to-mesenchymal transition, a hallmark of cancer progression (50–52). Given the array of stresses that induce SINE transcription, there are likely additional functions of SINE ncRNAs yet to be uncovered.

Genome-wide mapping studies in murine fibroblasts identified widespread induction of B2 SINEs and upregulation of ∼14% of tRNA loci upon MHV68 infection (45, 53). This induction initiates prior to viral genome replication and is not a consequence of antiviral signaling, nor does it involve Maf1, a central negative regulator of RNAPIII activity (53, 54). A partial screen of MHV68 open reading frames (ORFs) identified a role for the conserved herpesvirus kinase ORF36 in B2 SINE activation, potentially through alterations to the chromatin landscape (53, 54). However, MHV68 mutants lacking functional ORF36 still induce modest levels of SINE RNA (53, 54), suggesting that additional viral activities contribute to RNAPIII activation.

Here, we identify a second MHV68 protein, ORF45, as an activator of B2 SINE and pre-tRNA transcription during infection. We show that MHV68 ORF45 interacts with ORF36 and together these proteins additively increase RNAPIII transcription. Increased RNAPIII activity requires the ability of ORF45 to stimulate the extracellular signal-regulated kinase (ERK) mitogen-activated protein kinase (MAPK) pathway, which increases the levels of Brf1, an essential component of the RNAPIII transcription factor complex TFIIIB. We show that other activators of MAPK/ERK signaling also enhance Brf1 expression, suggesting that this is a common mechanism to increase RNAPIII activity under conditions where components of the RNAPIII transcriptional machinery are limiting.

## Materials & Methods

### Plasmids and cloning

MHV68 FLAG tagged ORF45 and ORF65 were subcloned into the XhoI and NotI sites of pcDNA4/TO-3xFLAG (N-terminal tag) to generate pcDNA4/TO-3xFLAG-ORF45 or ORF65 using InFusion cloning (Clontech). ORF45 and ICP0 was subcloned into the BamHI and XhoI sites of pcDNA4/TO-2xStrep (N-terminal tag) to generate pcDNA4/TO-2xStrep-ORF45 using InFusion cloning (Clontech). Deletion mutants of 2xStrep-ORF45 were generated using site-directed mutagenesis PCR with Q5 DNA Polymerase (New England Biolabs) with primers listed in Table 1. PCR products were DpnI treated, ligated using T4 polynucleotide kinase and T4 DNA ligase, and transformed into Escherichia coli XL-1 Blue cells. To generate pLKO.1-ORF45 shRNA-1 and pLKO.1-ORF45 shRNA-2 for lentivirus transductions, the pLKO.1-TRC cloning vector (Addgene plasmid 10879) was digested with EcoRI and AgeI to release a 1.9kb stuffer. shRNA oligos targeting ORF45 listed in Table 1 were then annealed and ligated into the vector using T4 DNA ligase (New England Biolabs) and transformed into Escherichia coli XL-1 Blue cells. The packaging plasmids pMD2.G (Addgene plasmid 12259) and psPAX2 (Addgene plasmid 12260) were also used to generate lentivirus. All newly generated plasmids will be deposited to Addgene prior to publication of the manuscript.

**Table 1.**
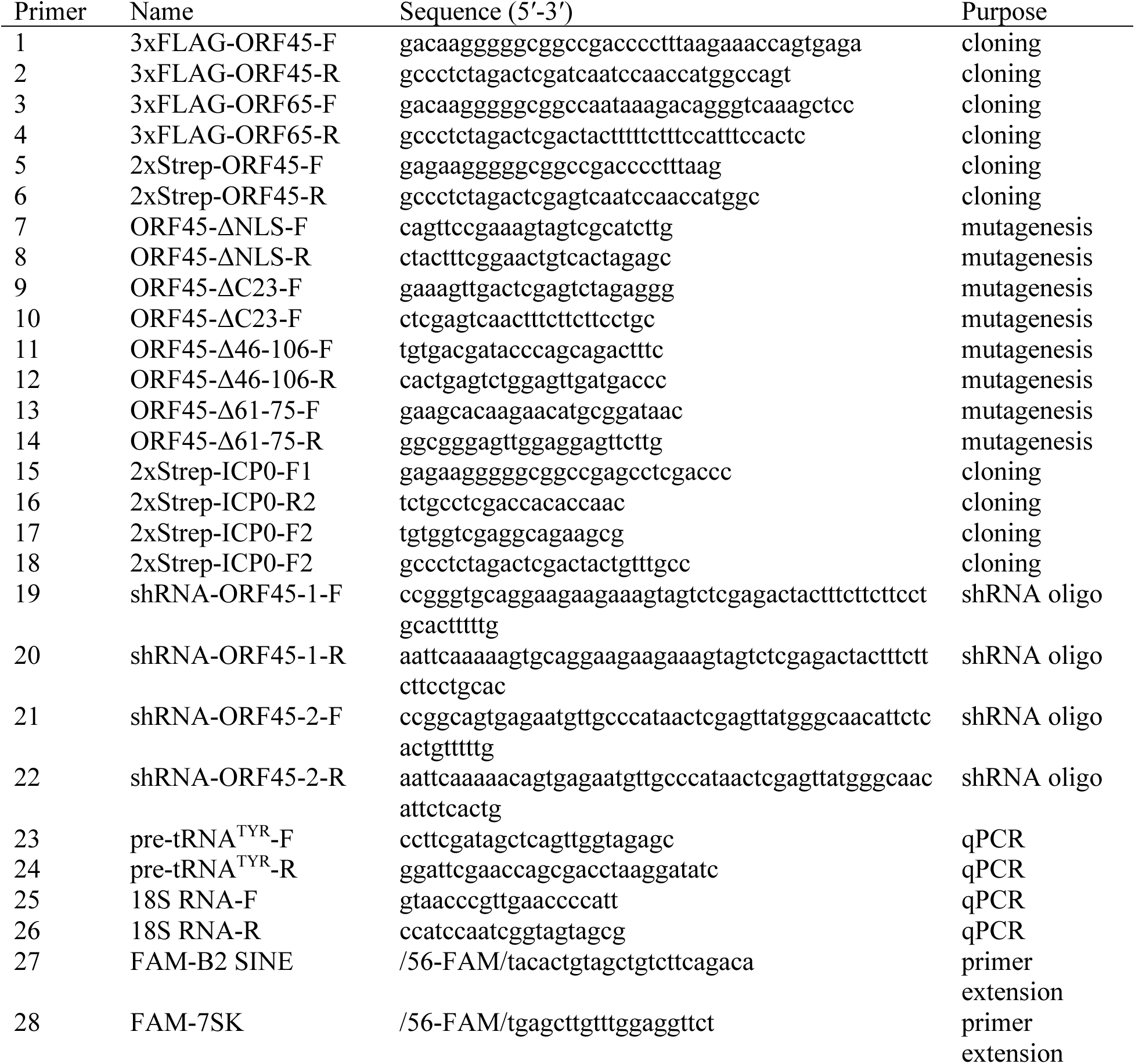
Primers used in this study

### Cell lines and transductions

NIH 3T3 (ATCC CRL-1658) and NIH 3T12 (ATCC CCL-164) mouse fibroblast cell lines and HEK293T human cell lines (ATCC CRL-3216) were maintained in Dulbecco’s modified Eagle’s medium (DMEM; Gibco) with 10% fetal calf serum (FBS; VWR) and screened regularly for mycoplasma by PCR. To generate transduced NIH 3T3 cell lines stably expressing a control shRNA or MHV68 ORF45 targeting shRNAs, lentivirus was generated in HEK293T cells by co-transfection of the pLKO.1 TRC cloning control vector or ORF45 expression shRNA vectors along with packaging plasmids. 24 h after transfection, the media was replaced with fresh DMEM supplemented with 10% FBS and 10μg/ml bovine serum albumin (BSA; Invitrogen). After 48 h, the supernatant was harvested, and syringe filtered through a 0.45-μm-pore-size filter (Millipore). Polybrene was added to a final concentration of 8 μg/ml to freshly trypsinized NIH 3T3 cells (3 × 106) and cells were spinoculated with 2ml of the lentivirus containing supernatant in a 12-well plate for 2 h at 1,000 × g. After 24 h, the cells were expanded to a 10-cm tissue culture plate and selected for 1 week in media supplemented with 2μg/ml puromycin (MilliporeSigma).

### Transfections

Transfections of MHV68 ORF containing plasmids into NIH 3T3 fibroblasts were completed as follows: 3T3 cells were maintained in DMEM with 10% FBS and grown to 90% confluency. Cells were removed and washed once with DPBS (Gibco). Transfections were done using the Neon Transfection System (Thermo Fisher). 2×10^6^ cells were resuspended in 100μl of buffer R, to which GFP (30 μg), ORF45 (30 μg), ORF65 (30 μg), ORF36 (5 or 10 μg), ICP0 (30 μg), or Ras-V12 (20 μg) plasmid DNA was added (in the cases where less that 30 μg of DNA was used, empty pcDNA4/TO-vector was added to equal 30 μg of total plasmid DNA). This was loaded into a Neon 100μl pipette tip and Neon tube with 3ml of buffer E2 with electroporation parameters set to 1300 V, 20 ms, 2 pulses. Following electroporation, cells were plated in 2ml of DMEM with 10% FBS in 6-well TC-treated plates and were incubated at 37C for 24 h.

### Virus preparations and infections

MHV68 was amplified in NIH 3T12 fibroblast cells, and the viral 50% tissue culture infective dose (TCID50) was measured on NIH 3T3 fibroblasts by limiting dilution. NIH 3T3 fibroblasts were infected at the indicated multiplicity of infection (MOI) by adding the required volume of virus to cells in 5 ml of serum free DMEM in 10cm TC-treated plates. Infection was allowed to proceed for 1 h to allow for viral entry followed by removal of the virus containing media and replacement with fresh DMEM with 10% FBS.

### Primer extension and RT-qPCR

Total RNA was extracted from cells using TRIzol reagent (Invitrogen). Primer extension was performed on 15μg of total RNA using a 5′-fluorescein labeled oligos specific to B2 SINE RNA or 7SK RNA. RNA was ethanol precipitated in 1 ml 100% EtOH, washed in 70% ethanol and pelleted at 21,130 × g and 4°C for 10 min. Pellets were resuspended in 18μl of 1X SuperScript III Reverse Transcriptase reaction buffer (SSIII-RT; Thermo Fisher) containing 1μl of each 5′-fluorescein labeled primer (10 pmol/μl) listed in Table 1. Samples were heated to 80°C for 10 min, followed by annealing for 1 h at 56°C. Then, 30μl of extension buffer (1X SSIII-RT buffer, 40U RNasin Ribonuclease Inhibitor (Promega) 2 mM DTT, 1 mM dNTP, 1000U of SSIII-RT) was added, and extension was carried out for 1 h at 42°C. Samples were precipitated in 100% ethanol for 20 min at -80°C, and then pellets were briefly air dried and resuspended in 20μl 1× RNA loading dye (47.5% formamide, 0.01% SDS, 0.01% bromophenol blue, 0.005% xylene cyanol, and 0.5 mM EDTA). Then, each sample was run on an 8% urea-PAGE gel for 1 h at 250 V. Gels were imaged on a Chemidoc imager (Bio-Rad) with fluorescein imaging capability. Relative induction of B2 SINE ncRNAs for each sample was measured as the ratio of the mean integrated intensity between 7SK RNA and B2 SINE ncRNA level using FIJI (55) and was normalized to the GFP-expressing plasmid control. For RT-qPCR, total RNA was isolated from cells using TRIzol (Invitrogen), treated with Turbo DNase (Ambion), and reverse transcribed with AMV RT (Promega) primed with random 9-mers. Quantitative PCR (qPCR) analysis was performed with iTaq Universal SYBR green Supermix (Bio-Rad) using the primers listed in Table 1. qPCR was performed on at least three biological replicates and threshold cycle (CT) values were measured from three technical replicates per biological sample. Fold change was calculated by ΔΔC_T_ method.

### Affinity purification and western blotting

To prepare whole cell lysates for evaluating protein expression of viral ORFs 24 h after transfection, cells were washed with cold DPBS (Gibco) followed by lysis with RIPA lysis buffer (50 mM Tris HCl, 150 mM NaCl, 1.0% [vol/vol] NP-40, 0.5% [wt/vol] sodium deoxycholate, 1.0 mM EDTA, and 0.1% [wt/vol] SDS, cOmplete, Mini, EDTA-free Protease Inhibitor Cocktail - Roche). Cell lysates were vortexed briefly, rotated at 4°C for 15 min, and then clarified by centrifugation at 21,000 × *g* in a tabletop centrifuge at 4°C for 10 min to remove debris. Cell lysates for affinity purification were prepared at 24 h after transfection by washing and pelleting cells in cold DPBS, followed by resuspension in lysis buffer (50 mM Tris HCl, 150 mM NaCl, 0.5% [vol/vol] NP-40, 1.0 mM EDTA, cOmplete, Mini, EDTA-free Protease Inhibitor Cocktail – Roche) and rotation at 4°C for 30 min. Lysates were clarified by centrifugation at 21,000 × g at 4°C for 10 min, and then 1mg of lysate was incubated with prewashed MagStrep “type 3” XT magnetic beads (IBA LifeSciences) overnight in IP buffer (150 mM NaCl, 50 mM Tris-HCl, pH 7.4). The beads were washed 3 times for 5 min each with IP wash buffer (150 mM NaCl, 50 mM Tris-HCl, pH 7.4, 0.05% NP-40) and eluted with 2× Laemmli sample buffer (Bio-Rad). 30 μg of whole-cell lysate or IP elutions were resolved on 4 to 15% mini-PROTEAN TGX gels (Bio-Rad). Transfers to PVDF membranes (Bio-Rad) were done with the Trans-Blot Turbo transfer system (Bio-Rad). Blots were incubated in 5% milk in TBS with 0.1% Tween 20 (TBS-T) to block, followed by incubation with primary antibodies against Strep Tag II Antibody HRP conjugate (Sigma, 1:5000), anti-FLAG Antibody (Sigma F7425, 1:2500), anti-GAPDH monoclonal antibody (Invitrogen AM4300, 1:5000), anti-Vinculin antibody (abcam ab91459, 1;5000), rabbit polyclonal antiserum to ORF45 (generously provided by Dr. Ren Sun (56)), anti-phospho-p44/42 MAPK (Erk1/2) (Thr202/Tyr204) antibody (Cell Signaling #9101, 1:1000), anti-Brf1polyclonal antibody (Bethyl, 1:1000), or anti Ha-Ras antibody (Sigma MC57, 1:1000) in 5% milk in TBS-T. Washes were carried out with TBS-T. Blots were then incubated with horseradish peroxidase (HRP)-conjugated secondary antibodies (Southern Biotechnology, 1:5000). Washed blots were incubated with Clarity Western ECL substrate (Bio-Rad) for 5 min and visualized with a ChemiDoc imager (Bio-Rad).

### Immunofluorescence

At 24h post-transfection, NIH 3T3 fibroblasts were plated on coverslips (7.5 x 10^4^ cells/well of a 12-well plate) then fixed in 4% formaldehyde for 10 min. Cells were permeabilized with ice-cold methanol at -20°C for 20 min and incubated with rabbit polyclonal antiserum to ORF45 (generously provided by Dr. Ren Sun (56); 1:200) in 5% BSA overnight at 4°C. Then Alexa Flour 488 goat anti-rabbit IgG antibody (Invitrogen) was added (1:1000) for 1 h at room temperature. Coverslips were mounted in DAPI-containing Vectashield (VectorLabs). Imaging was performed using a confocal Zeiss LSM 880 NLO AxioExaminer microscope driven by Zen 2 software with a 40x water immersion objective (Zeiss & 1.0 NA). Image analysis was performed in FIJI, including image cropping, and brightness/contrast adjustments (55).

## Results

### MHV68 ORF45 is sufficient to induce B2 SINE transcription and is required for robust B2 SINE and pre-tRNA transcription during MHV68 infection

Previously, we screened a partial library of MHV68 open reading frames (ORFs) and identified ORF36 as an activator of B2 SINE transcription (53). To determine if any additional MHV68 ORFs induce B2 SINE transcription, we cloned and re-screened a portion of the library that included an additional 19 FLAG tagged ORFs not represented in our prior screen. These were individually transfected into NIH 3T3 fibroblasts and B2 SINE ncRNA levels were measured by B2 SINE specific primer extension (data not shown). Among these, expression of only one additional MHV68 gene was sufficient to upregulate B2 SINEs. Cells expressing FLAG-ORF45, a virion tegument protein conserved among the *Gammaherpesvirinae*, showed a ∼2.5-fold increase in B2 SINE ncRNA levels compared to a GFP-expressing plasmid control (Fig. 1A). In line with prior observations that MHV68 specifically activates B2 SINE and pre-tRNA loci (39, 45), ORF45 did not alter the levels of another RNAPIII transcribed small nuclear RNA, 7SK. ORF45-mediated induction of B2 SINEs was less pronounced than the ∼8-fold induction of B2 by ORF36, perhaps due to the lower expression levels of FLAG-ORF45 compared to FLAG-ORF36. (Fig. 1B). Nonetheless, B2 SINE transcriptional activation was consistently specific to only ORF45 or ORF36 and not observed with other MHV68 ORFs such as ORF65 (Fig. 1A, data not shown).

**FIG 1.**
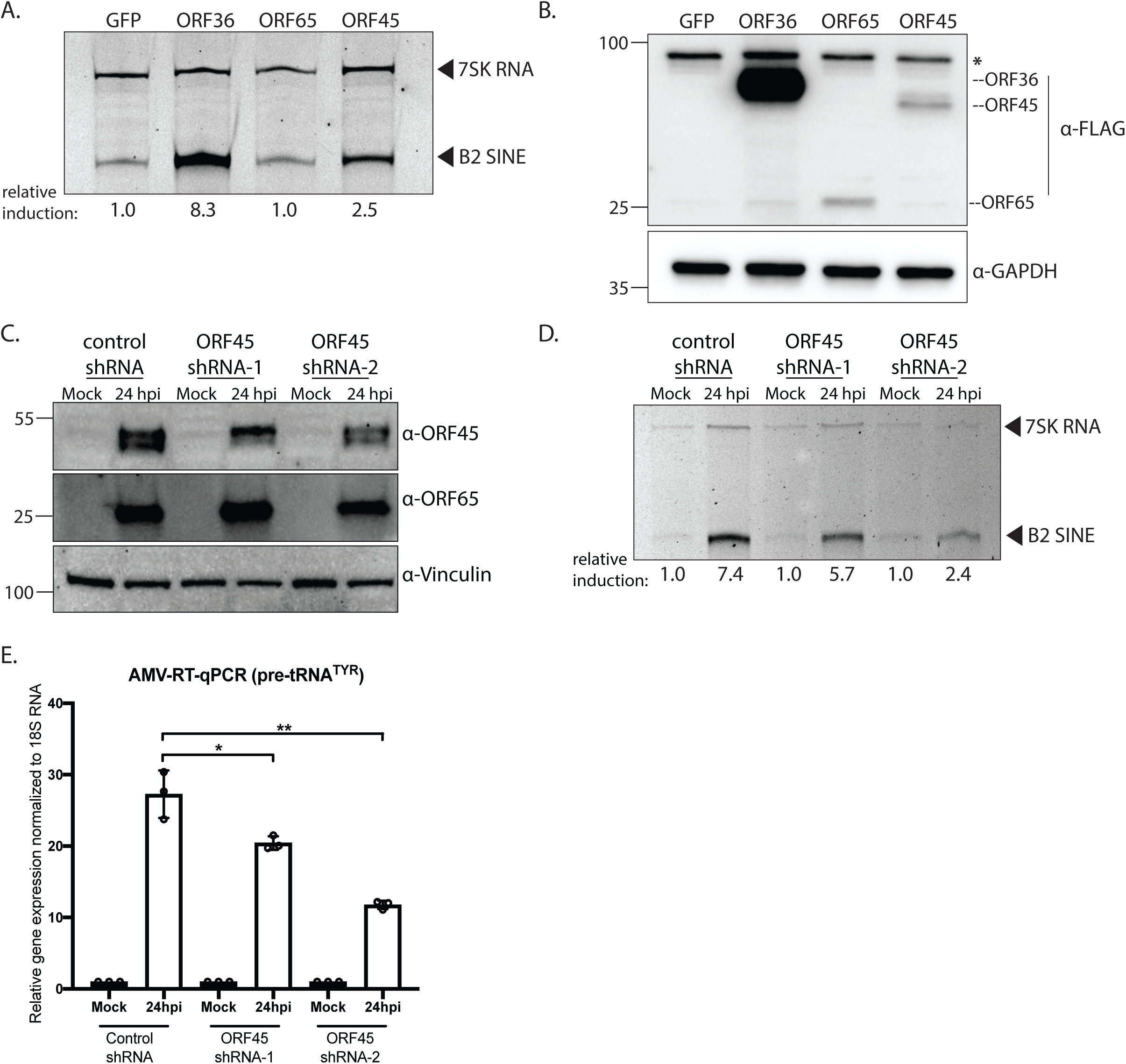
MHV68 ORF45 induces B2 SINE transcription in murine fibroblasts contributes to B2 SINE and pre-tRNA transcription during MHV68 lytic infection. (A) NIH 3T3 cells were transfected with plasmids containing the indicated N-terminally FLAG tagged MHV68 ORFs or GFP expressing control for 24 h, whereupon total RNA was subjected to primer extension using primers specific to B2 SINE ncRNAs or 7SK RNA (control). In this and all subsequent primer extensions, the relative induction of B2 SINE ncRNAs for each sample was measured as the ratio of the mean integrated intensity between 7SK RNA and B2 SINE ncRNA levels and normalized to the control (GFP-expressing plasmid) ratio. (B) Protein extracted from cells harvested from (*A*) was analyzed by Western blotting with antibodies against FLAG and GAPDH (loading control). *Represents a non-specific band. (C) Stable NIH 3T3 cell lines constitutively expressing a control shRNA or ORF45 targeting shRNAs (shRNA-1 or shRNA-2) were mock infected or infected with MHV68 at an MOI of 5. At 24hpi cells were harvested and lysed to extract total protein and was analyzed by Western blotting with antibodies against MHV68 ORF45, ORF65, and Vinculin (loading control). (D) Cells from (C) were also harvested for total RNA extraction and subjected to primer extension using primers specific to B2 SINE ncRNAs or 7SK RNA (control). (E) Total RNA extracted from (D) was also used to detect pre-tRNA^TYR^ levels with forward and reverse primers with 3′ ends complementary to the tRNA intron using AMV RT-qPCR. Expression was normalized to 18S rRNA and compared to values from mock-infected cells. RT-qPCR experiments were done in triplicate. Error bars show the standard deviation (SD), and statistics were calculated using an unpaired *t* test on raw Δ*C_T_* values. ns, not significant. *, *P* < 0.05 **, *P* < 0.01.

We next sought to assess the contribution of ORF45 towards B2 SINE and pre-tRNA transcriptional activation during infection. ORF45 is an essential MHV68 gene, so to deplete it we generated 3T3 fibroblast cell lines constitutively expressing control or ORF45 targeting shRNAs. The ORF45-targeting shRNAs partially reduced ORF45 protein expression 24 hours post infection (hpi) with MHV68 when compared to the control shRNA cell line, whereas expression of the ORF65 capsid protein was largely unaffected (Fig. 1C). Reduced ORF45 levels scaled with a reduction in MHV68-induced B2 RNA as measured by primer extension, and in pre-tRNA induction as measured by quantitative reverse transcriptase PCR (RT-qPCR) using primer sets that were specific for the pre-tRNAs produced from the intron-containing tRNA-TYR gene (Fig. 1D-E). From these data we conclude that ORF45 contributes to RNAPIII transcriptional activation during MHV68 infection.

### MHV68 ORF45 and ORF36 interact and additively upregulate B2 SINE and pre-tRNA transcription

In KSHV, the ORF45 and ORF36 proteins interact (57), which is notable given that these are the two MHV68 ORFs involved in RNAPIII activation. In streptavidin-based affinity purification experiments, we similarly found that N-terminally Strep tagged MHV68 ORF45 associated with with N-terminally FLAG tagged MHV68 ORF36 (Fig. 2A). Also, in line with studies of the KSHV protein homologs (57), expression of MHV68 ORF36 resulted in increased levels of the ORF45 protein (Fig. 2B). This did not appear to be a mutually stabilizing interaction, as ORF36 protein levels were unaffected by co-transfection of ORF45 (Fig. 2B).

**FIG 2.**
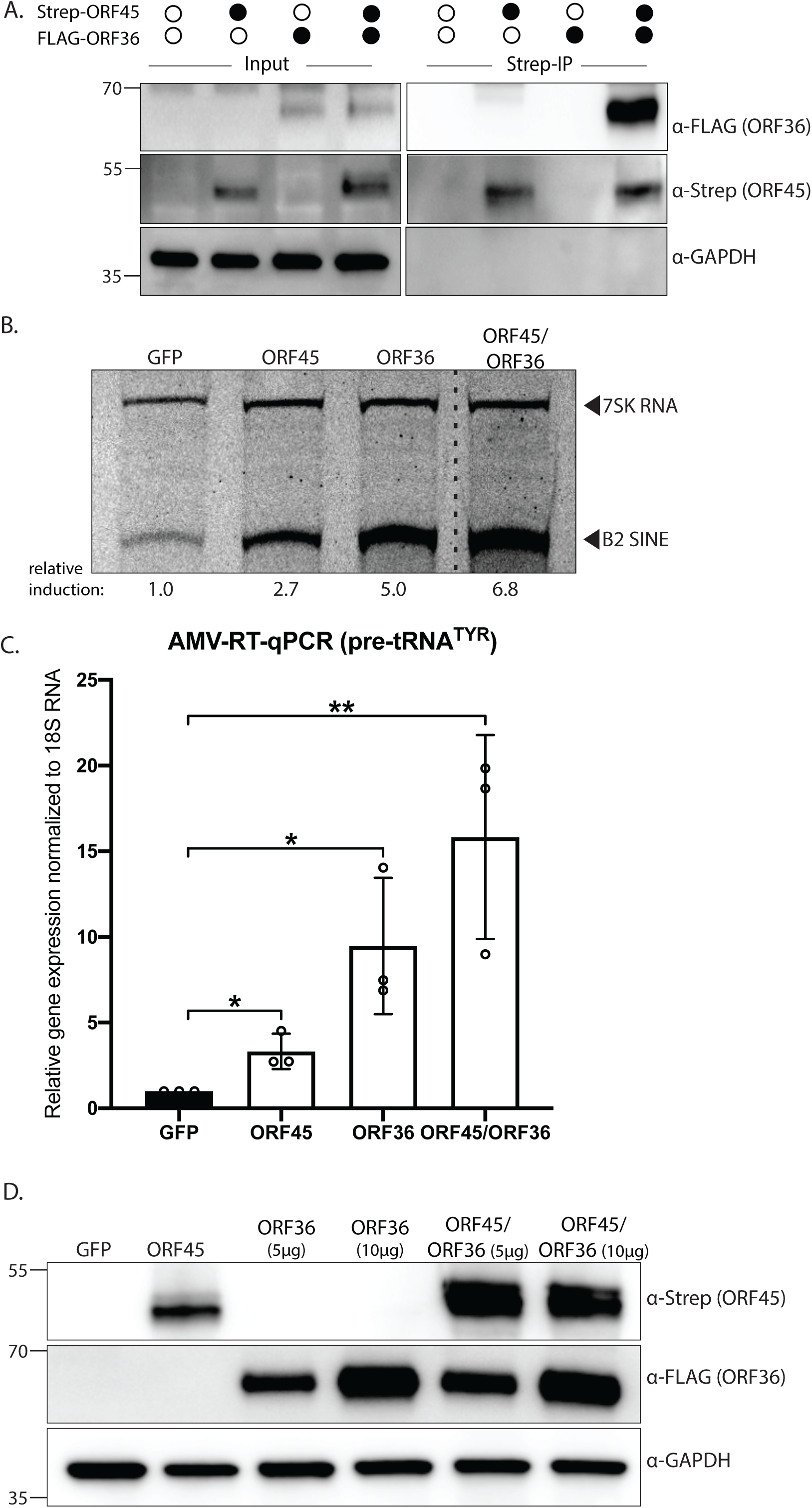
MHV68 ORF45 and ORF36 interact and mutually enhance B2 SINE and pre-tRNA transcription. (A) NIH 3T3 fibroblasts were transfected with Strep-ORF45 or FLAG-ORF36 and lysates were purified over StrepTactinXT magnetic beads followed by western blotting with antibodies against Strep, FLAG, and Vinculin (loading control). (B-D) NIH 3T3 fibroblasts were transfected with Strep-ORF45, FLAG-ORF36, or a GFP control for 24 h. Protein lysates were analyzed by Western blotting with antibodies against Strep, FLAG, and Vinculin (loading control) (B). RNA was subjected to either primer extension using primers specific to B2 SINE ncRNAs or 7SK RNA (as a control) (*C*), or RT-qPCR to detect pre-tRNA^TYR^ (D). In (C), the dashed line indicates where an irrelevant lane was removed from the image. In (D), pre-tRNA^TYR^ levels were normalized to 18S rRNA and compared to values from the GFP expressing control. RT-qPCR experiments were done in triplicate. Error bars show the standard deviation (SD), and statistics were calculated using an unpaired *t* test on raw Δ*C_T_* values. ns, not significant. *, *P* < 0.05 **, *P* < 0.01.

Given that both ORF45 and ORF36 can independently activate RNAPIII transcription and that and their interaction boosts ORF45 expression, we considered whether their interaction additively or synergistically impacted RNAPIII. Cells co-expressing Strep-ORF45 and FLAG-ORF36 displayed an additive increase of B2 SINE expression (∼7 fold) when compared to cells only expressing ORF45 (∼3 fold) or ORF36 (∼5 fold) (Fig. 2B). Similarly, pre-tRNA^TYR^ levels increased by ∼3 and ∼9 fold in the presence of ORF45 or ORF36, respectively, and additively increased to ∼15-fold upon their co-transfection (Fig 2C). Thus, although MHV68 ORFs 45 and 36 interact, their stimulation of RNAPIII transcription may occur by distinct mechanisms.

### The ORF45 NLS and conserved C-terminal domain are dispensable for B2 SINE induction

Both KSHV ORF45 and MHV68 ORF45 contain nuclear location signals and can traffic between the nucleus and the cytoplasm; however, KSHV ORF45 is predominantly localized to the cytoplasm while MHV68 ORF45 is predominantly localized to the nucleus (56, 58). Deletion of the MHV68 ORF45 nuclear localization (Strep-ORF45-ΔNLS) signal shifted its localization largely to the cytoplasm as measured by immunofluorescence assay (Fig. 3A). Despite this shift in localization, Strep-ORF45-ΔNLS activated B2 SINE transcription to comparable levels as WT ORF45, suggesting that a predominant nuclear localization may be dispensable for B2 SINE activation (Fig. 3B).

**FIG 3.**
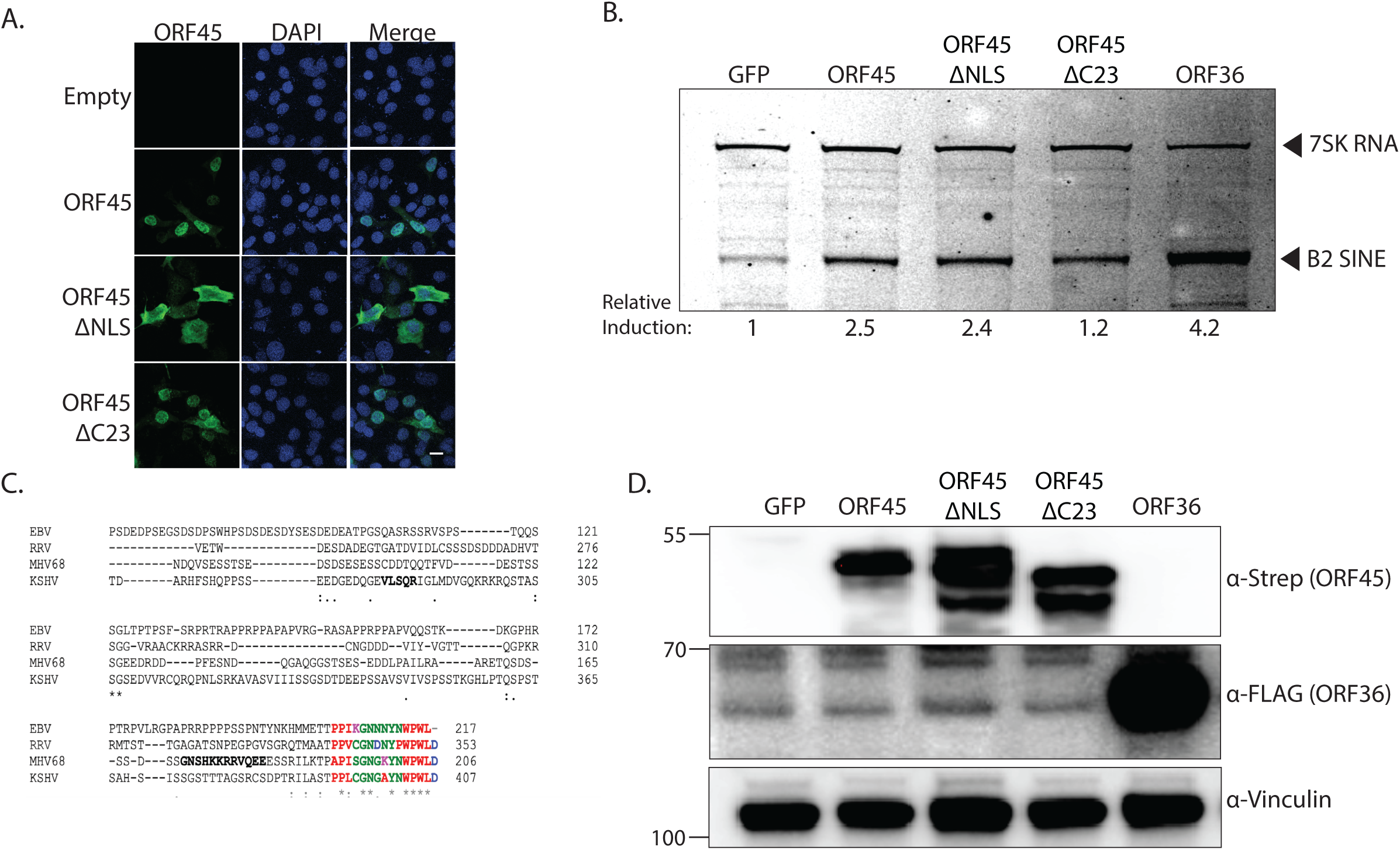
The ORF45 NLS and C-terminal residues are dispensable for B2 SINE transcriptional activation. (A) Confocal microscopy images of NIH 3T3 cells transfected with an empty control vector or the indicated strep tagged ORF45 plasmids and incubated with anti-Strep antibodies to detect ORF45 (green) or DAPI to identify nuclei (blue), with the right panels showing merged images. Scale bar = 5 µm. (B) NIH 3T3 cells were transfected with plasmids containing indicated Strep tagged genes or a GFP control for 24 h, whereupon cells were harvested for total RNA extraction and subjected to primer extension using primers specific to B2 SINE ncRNAs or 7SK RNA (control). (C) Sequence alignment of ORF45 homologs. Diagram shows the most conserved regions of KSHV ORF45 (amino acids 305-407), EBV BKRF4 (amino acids 121-217), MHV68 ORF45 (amino acids 122-206), and RRV ORF45 (276-253). NLS sequences in KSHV and MHV68 ORF45 are bolded, and the C-terminal regions are highlighted with color to show the most conserved residues (D) Protein lysates from cells harvested from (*B*) were analyzed by Western blotting with antibodies against Strep, FLAG, and Vinculin (loading control).

Gammaherpesvirus ORF45 homologs share very low amino acid sequence identity, although the C-terminal region is the most conserved and is required for transcomplementation of a MHV68 ORF45-null mutant virus (56). The last 23 amino acids of the MHV68 ORF45 C-terminus share 58% identity to the C-terminus of KSHV ORF45, 50% to the EBV homolog BKRF4, and 39% to the Rhesus rhadinovirus (RRV) ORF45 homolog (Fig. 3C). However, deletion of the C-terminal 23 amino acids (ORF45-ΔC23) resulted in only a modest reduction of B2 SINE transcriptional activation (Fig. 3B), although all three ORF45 proteins are comparably expressed and exhibited the characteristic protein doublet indicative of phosphorylation (Fig. 3D). Thus, neither strong nuclear localization nor the conserved C-terminal region of MHV68 ORF45 are essential for RNAPIII activation.

### Upregulation of RNAPIII activity by MHV68 ORF45 requires stimulation of MAPK/ERK signaling

KSHV ORF45 interacts with the cellular p90 ribosomal S6 kinase (RSK1/2) and forms a ternary complex with the extracellular signal-regulated kinase (ERK2) of the mitogen-activated protein kinase (MAPK) pathway, strongly stimulating their kinase activities (59–65). This correlates with sustained phosphorylation of RSK1/2 and ERK1/2, which are functional mediators of MAPK/ERK signaling, and are important for MHV68 lytic infection (66). In agreement with data for KSHV ORF45, transfection of MHV68 Strep-ORF45 greatly increased the abundance of phosphorylated (i.e., active) ERK1/2, indicative of activated MAPK/ERK signaling (Fig 4A, B). Amino acids 46-106 of MHV68 ORF45 are most similar to the previously identified RSK/ERK activation domain of KSHV ORF45 (67) (Fig 4A). Expression of an ORF45 deletion mutant lacking this region (Strep-ORF45-Δ46-106) resulted in reduced levels of phosphorylated ERK1/2, however the deletion also impaired ORF45 protein expression, confounding interpretation of its activity (Fig 4B). We tested additional mutants and ultimately determined that a smaller deletion of amino acids 61-75 (ORF45-Δ61-75) resulted in protein expression levels comparable to the full-length protein yet did not activate ERK1/2 (Fig. 4C & data not shown). Unlike full length ORF45 or the ORF36 control, ORF45-Δ61-75 failed to induce B2 SINE RNA as measured by primer extension (Fig 4D). Overall, these data validate a role for MHV68 ORF45 in the activation of cellular MAPK/ERK signaling, identify an ERK activation domain within MHV68 ORF45, and demonstrate a connection between ORF45 mediated ERK MAPK signaling and RNAP III transcriptional activation.

**FIG 4.**
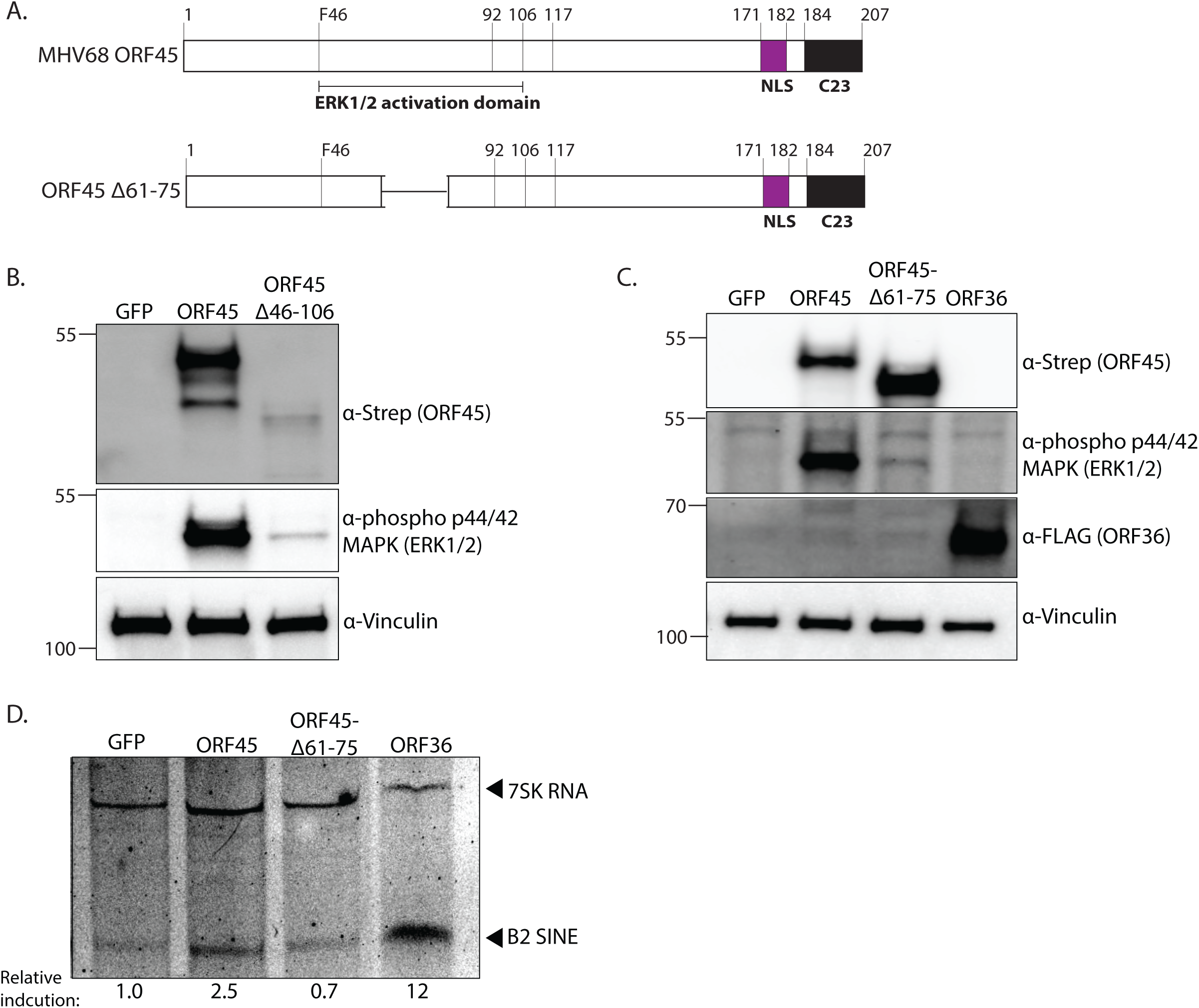
MHV68 ORF45-induced MAPK/ERK signaling is required for B2 SINE and pre-tRNA induction. (A) Schematic of MHV68 ORF45 protein showing the putative ERK1/2 activation domain. (B-C) NIH 3T3 fibroblasts were transfected with indicated Strep tagged genes or a GFP control for 24h, whereupon lysates were analyzed by Western blotting with antibodies against Strep, phospho-ERK1/2, FLAG, and Vinculin (loading control). (D) NIH 3T3 fibroblasts were transfected with Strep tagged ORF45, ORF45Δ61-75, or a GFP control for 24h, whereupon levels of B2 SINE or 7SK (control) RNA were determined by primer extension. Cells were also lysed to extract total protein and was analyzed by Western blotting (right) with antibodies against Strep, phospho-ERK1/2 and Vinculin (loading control).

### Constitutive activation of MAPK/ERK signaling leads to B2 SINE and pre-tRNA upregulation through increased levels of Brf1

Several connections exist between RNAPIII activity and mitogenic factors such as ERK. Most notably, ERK directly binds and activates components of the essential RNAPIII transcription factor complex, TFIIIB, and the levels of the essential TFIIIB component Brf1 increase in response to cellular growth stimuli in specific cell types (68, 69) (70, 71). We therefore hypothesized that ORF45 might increase RNAPIII activity by boosting the levels of Brf1.

Indeed, while endogenous Brf1 protein was barely detectable in NIH 3T3 fibroblasts transfected with a control GFP plasmid, its levels markedly increased upon transfection of ORF45 (Fig 5A). This was dependent on activation of ERK signaling, as Brf1 levels did not increase in cells containing the ORF45-Δ61-75 mutant (Fig 5A). Also notable was that MHV68 ORF36 did not alter Brf1 levels, indicating that its RNAPIII induction occurs via a mechanism distinct from that of ORF45 (Fig 5A).

**FIG 5.**
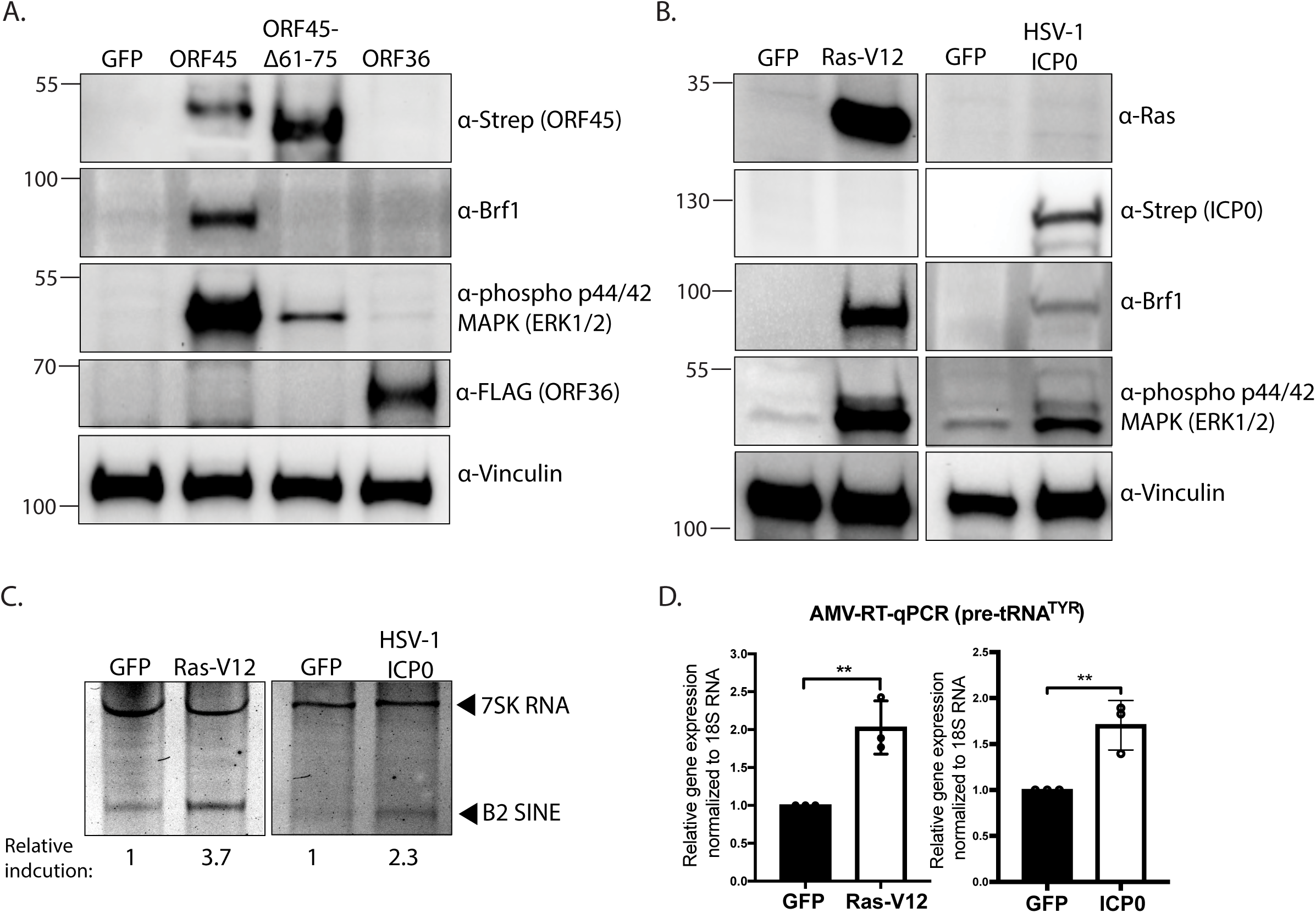
Constitutive activation of MAPK/ERK signaling upregulates Brf1 and enhances RNAPIII transcription. NIH 3T3 fibroblasts were transfected with the indicated expression plasmids for 24h. (A-B) Protein lysates were analyzed by Western blotting with the indicated antibodies. Vinculin served as a loading control. (C-D) Total RNA was also extracted from the samples in (*B*) and subjected to primer extension using primers specific to B2 SINE ncRNAs or 7SK RNA (as a control) (*C*) or RT-qPCR to detect pre-tRNA^TYR^ levels, whose expression was normalized to 18S rRNA and compared to values from the GFP expressing control. RT-qPCR experiments were done in triplicate. Error bars show the standard deviation (SD), and statistics were calculated using an unpaired *t* test on raw Δ*C_T_* values. ns, not significant. *, *P* < 0.05 **, *P* < 0.01.

Finally, to determine whether ERK activation was sufficient to upregulate Brf1 and induce transcription of B2 SINEs and pre-tRNA, we activated ERK signaling through two ORF45-independent mechanisms. We expressed either a constitutively active form of the small GTPase H-Ras (Ras-V12) (72), or the HSV-1 ICP0 protein, which is also an activator of MAPK/ERK signaling (73). Indeed, compared to the GFP control plasmid, both Ras-V12 and HSV-1 ICP0 transfection resulted in an increase in Brf1 protein levels (Fig 5B), as well as enhanced expression of B2 SINE ncRNAs and pre-tRNA^TYR^ levels (Fig 5C, D). These levels were comparable to the levels of transcriptional enhancement induced by ORF45 (Fig 2B, C). Collectively, these data support a model whereby the ERK activation function of ORF45 increases RNAPIII transcription by elevating the levels of Brf1 in cells where its expression is limiting. This mechanism likely extends to other viral activators of MAPK/ERK signaling that regulate RNAPIII activity during viral infection.

## Discussion

RNAPIII transcription is modulated during infection with several DNA viruses, leading to the upregulation of specific RNAPIII transcripts alteration of viral gene expression (39-43, 45, 46, 54, 74). Together with our prior work (53), we have now screened 90% of the known MHV68 ORFs (75) and identified two, ORF36 and ORF45, as having independent RNAPIII activation functions. Both MHV68 ORF36 and ORF45 are required for efficient viral replication and viral gene expression and ORF45 is also involved in virion morphogenesis (56, 76, 77). ORF36 and ORF45 are early genes, consistent with the viral DNA replication-independent early kinetics of B2 SINE and pre-tRNA transcriptional activation (39, 45). Although mechanistic details of how ORF36 activates RNAPIII transcription remain unknown, it is proposed to function by inhibiting proteins involved in the maintenance of a repressive chromatin landscape (53). Our mutational analysis of MHV68 ORF45 revealed that its stimulation of RNAPIII transcription is coordinated in the cytoplasm through its ability to activate MAPK/ERK signaling. Notably, while the impact of ORF36 and ORF45 on RNAPIII activation is additive and thus likely to occur via distinct mechanisms, these proteins physically interact, underscoring their potential coordination during infection. Altogether, these data provide novel insight into the function of ORF45 and link cellular MAPK signaling to RNAPIII transcriptional activity in the context of viral infection.

Our data suggest that MHV68 ORF45 displays functional conservation with KSHV ORF45, despite sharing little sequence identity. Like their KSHV homologs, we show here that MHV68 ORF45 and ORF36 stably interact (57). In KSHV, ORF36 is a substrate of ORF45-activated kinases and ORF45 enhances the kinase activity of ORF36; although untested, this may similarly occur for the MHV68 homologs. KSHV ORF45 causes sustained activation of p90 ribosomal kinases (RSKs) and extracellular regulated kinases (ERKs) (59–64), and subsequent studies also identified that KSHV ORF45, ORF36, and RSK form a stable complex (57). Another function of KSHV ORF45 is suppression of interferon regulatory factor 7 (IRF7), a crucial regulator of type I interferon gene expression (78–80). Given that both IRF3 and IRF7 are dispensable for B2 SINE transcriptional activation during MHV68 infection (53), if MHV68 ORF45 retains this function we hypothesize it would not be linked to RNAPIII activity.

Studies in proliferating mammalian cells have been critical in understanding the connection between cellular regulators of proliferation (e.g., ERK1/2) and RNAPIII transcriptional regulation. Upon mitogenic stimulation, cell growth is accompanied by ERK activation and a rapid increase in RNAPIII transcription (81–83). Activated ERK2 induces RNAPIII transcription through phosphorylation of Brf1, an essential component of the RNAPIII transcription factor TFIIIB (70). Phosphorylation of Brf1 enhances promoter recruitment of TFIIIB and RNAPIII, increasing the transcriptional output of RNAPIII. Brf1 can be limiting for the type 2 RNAPIII promoters found in tRNAs and B2 SINEs, with several cell types exhibiting low basal levels of Brf1 (68, 70, 84, 85). Indeed, we observed very low basal Brf1 levels in murine fibroblasts and found that ORF45-induced ERK activation led to an increase in Brf1 protein expression. We hypothesize that this may be a viral strategy to overcome limiting levels of Brf1, thereby facilitating RNAPIII activation in MHV68 infected cells.

Many DNA and RNA viruses hijack the MAPK/ERK signaling cascade to mediate viral internalization, dysregulate the cell cycle, regulate viral replication, and prevent cell death (86–89). MAPK/ERK signaling promotes viral reactivation from latency during KSHV infection, viral replication during lytic MHV68 infection, and the production of infectious progeny during both MHV68 and KSHV infection (66, 90–98). The pro-viral nature of MAPK/ERK activation is consistent with findings that B2 SINE ncRNAs enhance viral gene replication and expression (39). ORF45 is only conserved among gammaherpesviruses, although the alphaherpesvirus HSV-1 strongly induces RNAPIII transcription of endogenous human SINE elements (Alus) (37). Here, we show one example of an HSV-1 viral protein (ICP0) that has been previously linked to ERK activity during infection (73), activates ERK signaling, increases Brf1 protein levels, and enhances B2 SINE and pre-tRNA transcription upon transfection in murine fibroblasts. Interestingly ICP4, which has been identified to play a role in RNAPIII activation during HSV-1 infection, functionally interacts with ICP0 (37, 99). It is possible that there are additional viral activators of MAPK/ERK signaling, whose characterization should yield new insights into regulation of noncoding RNA production during infection.

## Acknowledgements

We thank all members of the Glaunsinger and Coscoy labs for their helpful suggestions and discussion. We thank Dr. Ren Sun (University of California, Los Angeles) for generously providing MHV68 ORF45 and ORF65 antibodies, and Allison Didychuk for cloning FLAG tagged MHV68 ORF45 & ORF65. This work is funded by R01CA136367 to BAG, who is also an investigator of the Howard Hughes Medical Institute.

